# Photosynthetically active radiation is required for seedling growth promotion by volcanic dacitic tuff breccia (Azomite)

**DOI:** 10.1101/2022.10.03.510703

**Authors:** Kent F. McCue, Elijah Mehlferber, Robert Reed, Alexis Ortiz, Jon Ferrel, Rajnish Khanna

## Abstract

A plant’s growth and development are shaped by its genome and the capacity to negotiate its environment for access to light, water, and nutrients. There is a vital need to understand the interactions between the plant, its physical environment, and the fertilizers used in agriculture. In this study, a commercially available volcanic ash fertilizer, Azomite^®^, characterized as dacitic (rhyolitic) tuff breccia (DTB), was tested for its effect on promoting early seedling vigor. Early growth and photomorphogenesis processes are well studied in Arabidopsis. Seedling assays under different light conditions were used to dissect the underlying mechanisms involved. These assays are well established and can be translated to agriculturally important crop plants. The volcanic ash fertilizer was tested at different concentrations on seedlings grown on basic media lacking sucrose either in continuous darkness (Dc), continuous Red (Rc), Far-Red (FRc), or White Light (WLc). Micronutrients in the volcanic ash significantly increased seedling growth under Rc and WLc, but not under Dc and FRc, indicating that photosynthetically active radiation (PAR) was required for the observed growth increase. Furthermore, red-light photoreceptor mutant, *phyB-9* lacked the growth response, and higher amount of fertilizer reduced growth in all conditions tested. These data suggest that light triggers the ability of the seedling to utilize micronutrients in volcanic ash in a dose-dependent manner. The methods described here can be used to establish mechanisms of activity of various nutrient inputs, and coupled with whole-genome expression profiling, can lead to better insights into optimizing nutrient field applications to improve crop production.

## Introduction

In flowering plants, the mature embryo is surrounded by seed to protect and support its survival. Seeds remain dormant until the conditions become favorable for germination and provide nutrients to support early seedling growth (Koornneef and Karssen, 1994). Seedlings growing in the subterranean darkness of soil undergo skotomorphogenesis or etiolation, a process distinct from seedling development in light or photomorphogenesis (Von Arnim and Deng, 1996). Etiolation is achieved by cell elongation in the hypocotyl, while the folded shoot meristem is protected by the apical hook curvature as the seedling grows through the soil to reach sufficient light for photoautotrophic growth (Von Arnim and Deng, 1996). This mechanism is conserved across vascular plants to prevent photooxidative damage, which is phototoxic to the seedling caused by premature light exposure of protochlorophyllide, a precursor of chlorophyll (Reinbothe et al., 1996).

Exposure to light triggers photomorphogenesis, which is marked by the inhibition of hypocotyl elongation, opening of the apical hook, cotyledon unfolding and initiation of chlorophyll biosynthesis (greening) (Quail, 2002a). During this process the seedling exhibits precisely orchestrated photosensitivity to optimize its growth and development under the incumbent conditions. This is achieved through the perception of light as a signal, which carries information about color, direction, intensity, and duration of light. In *Arabidopsis*, the magnitude of light inhibition of hypocotyl length is synchronized with the reciprocal increase in cotyledon area during photomorphogenesis (Quail, 2002a). Previously, this correlation has been used as a diagnostic criterion to reveal that only a small number of early light-regulated genes are necessary for optimal de-etiolation (Khanna et al., 2006). The relationship between hypocotyl length and cotyledon area in the wild type seedlings was set to unity under set light conditions, and 32 genes categorized as early light responsive from microarray profiles were selected for analysis using corresponding gene-mutant lines (Khanna et al., 2006). Only 7 out of 32 gene lesions resulted in aberrant de-etiolation phenotypes, characterized by strict reciprocal correlations exhibiting increased hypocotyl length and reduced cotyledon area (hyposensitive to light), or inhibited hypocotyl length with enlarged cotyledon area (hypersensitive to light), relative to the wild type under the given light conditions (Khanna et al., 2006). These studies identified new genetic lesions that disrupted normal progression of photomorphogenesis. About half of the remaining mutant lines displayed no significant phenotypes. These genes were determined to be responsive to light but played either redundant or indirect roles in early seedling development. Whereas, several other mutant lines exhibited three different responses, either parallel inhibition or parallel enhancement of cell expansion in both hypocotyl and cotyledon, or only hypocotyl inhibition with no significant effect on cotyledon area (Khanna et al., 2006). This latter set of genes are more likely to be affected in general cell growth processes either in one or both organs but are not directly involved in the process of photomorphogenesis.

During photomorphogenesis, early seedling establishment is subjected to other environmental factors, including temperature and the availability of water and nutrients. A complex intersection of light responses with responsiveness to other environmental conditions collectively shapes the seedling’s growth in the soil. Sensitivity to informational light signal is achieved through a collection of photoreceptors with some overlapping and some distinct functions. *Arabidopsis* seedlings monitor light wavelengths through multiple photoreceptor systems, which include five phytochromes (phyA-phyE) for red/far-red (R/FR) and some degree of blue (B) light, along with cryptochromes (cry1 and cry2), the Zeitlupe (ZTL) family and phototropins (phot1 and phot2) for blue/UV-A, and UV Resistance Locus 8 (UVR8) for UV-B light (Briggs and Huala, 1999; Briggs and Christie, 2002; Quail, 2002a; Quail, 2002b; Chen et al., 2004; Rizzini et al., 2011). Plants can sense light radiation ranging from UV-B to FR (280-800 nm), which includes photosynthetically active radiation (PAR, 400-700 nm) (McCree, 1973). Photosynthetic efficiency is highest under red photons (600-700 nm), followed by green (500-600 nm) and blue (400-500 nm) photons (Inada, 1976), and it drops rapidly under wavelengths longer than 685 nm extending into the FR range, known as the “red drop” (Emerson and Lewis, 1943). However, FR and PAR photons can act synergistically under simultaneous irradiation (Emerson et al., 1957; Zhen et al., 2021, 2022). As the seedling becomes photoautotrophic, its rate of growth is shaped by the degree of photosynthesis and transpiration under daily variations in temperature, humidity, and soil composition.

In addition to light, early seedling establishment is highly sensitive to concentrations of cellular metabolites such as sugars and amino acids. Sugars provide intermediary source of metabolism and act as signaling molecules (Rolland et al., 2006). In the dark, the embryo transitions from mitotic growth to differentiation and cell expansion, marked by increased sucrose uptake derived from the stored neutral lipids and triacylglycerols (Pyc et al., 2017), accompanied by changes in gene expression and the activities of metabolic enzymes (Weber et al., 2005). Seedling growth in darkness through the soil is further supported by a sucrose feedback loop to increase sucrose levels in cotyledons (Mu et al., 2022). Cotyledon greening is triggered by light through the suppression of Constitutively Photomorphogenic 1 (COP1), and it is facilitated by Phytochrome Interacting Factors (PIFs) through the suppression of protochlorophyllide accumulation in darkness (Duek and Fankhauser, 2005; Zhong et al., 2009). FR light inhibits greening by suppressing the light-dependent conversion of protochlorophyllide A to chlorophyllide A, which is required for chlorophyll biosynthesis (Barnes et al., 1996). Exogenous application of sucrose can remove this FR block of cotyledon greening (Barnes et al., 1996), suggesting that sugars promote greening. Concomitant assembly of chlorophylls along with proteins and lipids leads to thylakoid and chloroplast differentiation (Jarvis and Lopez-Juez, 2013). Chloroplast development can be divided into two distinct phases, namely Structure Establishment and Chloroplast Proliferation (Pipitone et al., 2021). These stages of development rely upon several integral components, such as the insertion of Mg^2+^ into protoporphyrin IX by Mg^2+^-chelatase, a two-step process that is tightly regulated by light, the circadian clock, and the photosynthetic electron transport, involving regulation at transcriptional and post-transcriptional levels (Masuda, 2008).

The switch from cell expansion growth in darkness to early seedling establishment and subsequent growth is achieved through a continuum of signaling and resource management. The underlying molecular components of photomorphogenesis are well studied and described to be regulated at the levels of transcriptome and proteome in response to environmental cues. Knowledge of these mechanisms can be applied to dissect molecular pathways related to plant fertilizers. We tested a highly mineralized complex silica ore volcanic ash, surface mined in Utah, Azomite (AZOMITE^®^ Mineral Products, Inc.) because of its sustainability potential as a fertilizer sourced from a naturally occurring volcanic deposit. In a previous study, it was found that Azomite volcanic ash (DTB) increased tomato fruit production and concurrently it modified the rhizosphere and root endosphere microbial community architecture (Mehlferber et al., 2022b). We hypothesized that the temporal changes in microbiome community structures were primarily driven by the nutrient substrates available to the microbial communities, and that the observed shift suggested an increase in higher carbon (from C2 to C6) compounds in the host plant endosphere and possibly root exudates (Mehlferber et al., 2022b). These results suggested that Azomite treatment somehow modified host plant nutrient profile thereby causing a microbiome shift. Further, these changes pointed towards a plausible increase in the accumulation of photosynthesis products. Based on these findings, we wanted to test whether micronutrients in form of DBT impact plant growth directly in a nutrient-free medium. To investigate this possibility, we utilized a well-established Arabidopsis seedling photomorphogenesis test for two reasons; one to unravel the mechanism of action, and secondly to develop a reliable bioassay for any similar fertilizer. Here we show that Azomite promotes seedling growth in *Arabidopsis* during photomorphogenesis in a dose-dependent manner, and its activity is PAR-dependent. Deeper understanding of these mechanisms is essential for the development of safe and environmentally sustainable fertilizers.

## Results

Arabidopsis seedlings grown for 4-days on ½ MS media plates without sucrose, supplemented with indicated amounts of ultrafine grade of Azomite, in Dc, Rc, or WLc are shown (Fig. 1). The initial light treatments were selected to assess any observable phenotypic differences in response to different concentrations, which were chosen based on relative amounts used in our greenhouse tomato trials. There was a significant increase in hypocotyl length (20%) in response to 0.5 g/L of Azomite in Rc and 0.1 g/L in WLc (Fig. 2). Higher concentrations under all conditions tested, including Dc, Rc, and WLc led to relatively reduced hypocotyl growth (Fig. 2). These experiments have been repeated several times with similar results.

**Fig. 1.**
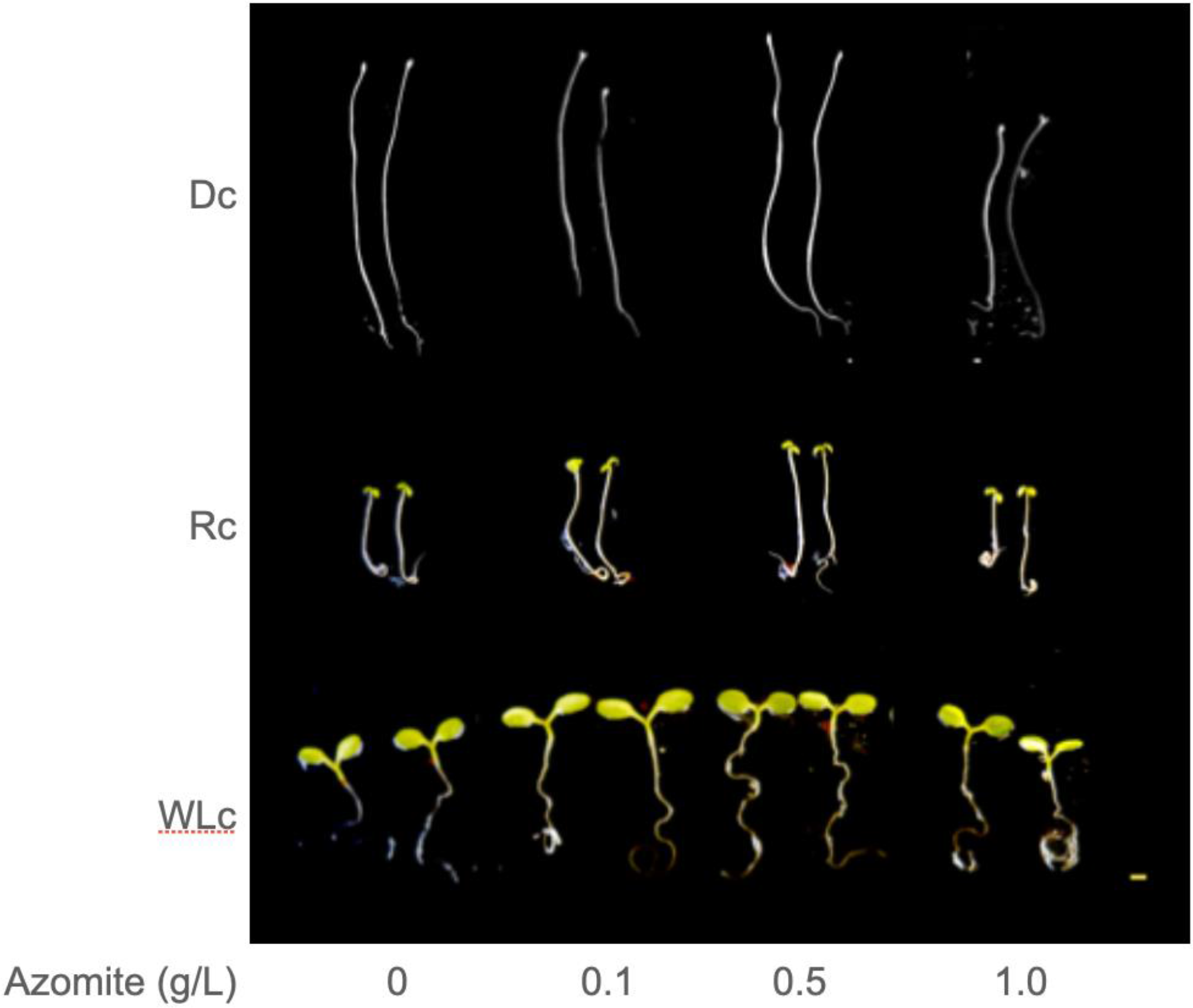
Arabidopsis seedlings respond differentially to increasing Azomite concentrations under light or darkness. Representative images of Col-0 seedlings grown for 4-days in continuous Darkness (Dc), Red light (Rc), or White Light (WLc) on media plates without sucrose, supplemented with different concentration of Azomite ultrafine (volcanic ash) as indicated.

**Fig. 2.**
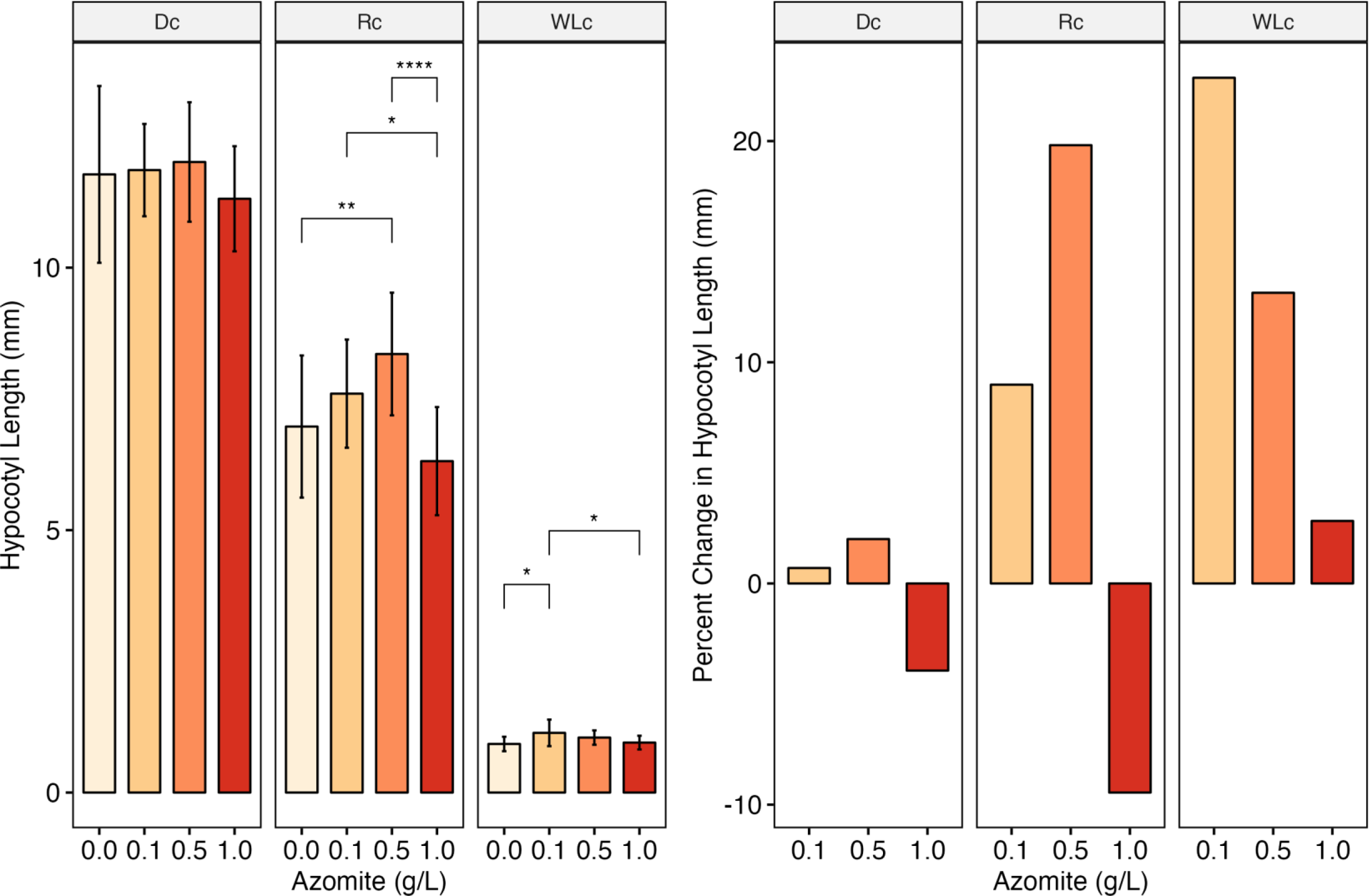
Azomite increases hypocotyl length under red and white light. Mean values of hypocotyl length of 4-day-old Col-0 seedlings in response to treatments with indicated Azomite concentrations grown in Dc, Rc or WLc (left). Percent change in hypocotyl length is shown for Azomite-treated seedlings, relative to untreated (0g/L) control (right). SE are plotted. P-values adjusted for multiple testing comparisons are represented as asterisks, with * indicating a p-value less than 0.05, ** indicating a p-value less than 0.01, and **** indicating a p-value less than 0.0001.

### Azomite micronutrients promote early seedling growth

Cotyledon area measurements revealed a similar pattern as seen with hypocotyl length, indicating that Azomite induced changes in cotyledon development under Rc and WLc. There was a significant increase in cotyledon area, 20% in Rc with 0.5g/L, and 25% in WLc with 0.1 g/L Azomite, and this effect on cotyledon area was reduced under higher concentration tested (Fig. 3). These initial results indicated that Azomite increased both hypocotyl elongation and cotyledon area under Rc and WLc in a dose-dependent manner (Figs. 2 and 3), but there was no significant effect on etiolated seedlings (Fig. 2). In addition, the peak increase in seedling growth under WLc required lower amounts of Azomite compared to the peak increase in growth under Rc (Figs. 2 and 3).

**Fig. 3.**
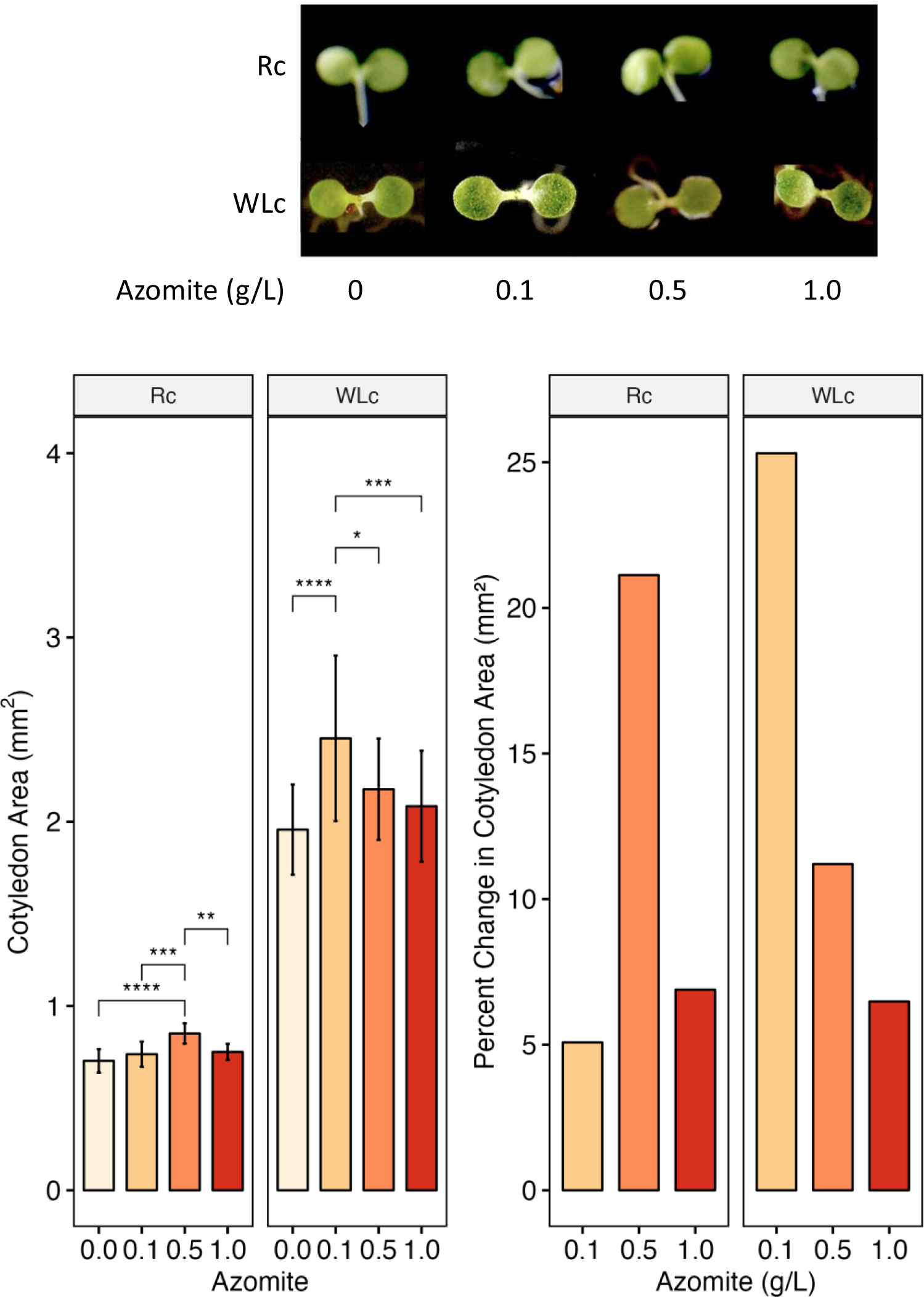
Azomite increases cotyledon area under red and white light. Representative images of cotyledons of 4-d-old Col-0 seedlings are shown in response to treatment with indicated Azomite concentrations under Rc and WLc (top). Mean values of cotyledon area of seedlings treated with different Azomite concentrations are plotted (left). Percent change in cotyledon area in response to Azomite treatment, relative to untreated (0g/L) control (right). SE are plotted. P-values adjusted for multiple testing comparisons are represented as asterisks, with * indicating a p-value less than 0.05, ** indicating a p-value less than 0.01, *** indicating a p-value less than 0.001, and **** indicating a p-value less than 0.0001.

Next, we tested photoreceptor *phyA-211* and *phyB-9* mutants under Rc and continuous Far-Red (FRc) light with Azomite treatments to test whether photosensitivity to different light wavelengths was linked to the observed response. As expected, *phyA-211* seedlings were insensitive to FRc, *phyB-9* seedlings showed reduced Rc sensitivity and Col-0 responded normally to the various light treatments (Fig. 4). There was no significant effect of Azomite treatments under Dc or FRc conditions on any of the genotypes tested (Fig. 5). There were significant changes in hypocotyl length elongation in Col-0 and *phyA-211* mutants in response to 0.5 g/L Azomite under Rc, exhibiting up to 20% and 10% increase in seedling growth, respectively (Fig. 5). These data indicated that photosensitivity to red light was required, and that Far-red or dark conditions were insufficient for Azomite activity on seedling growth.

**Fig. 4.**
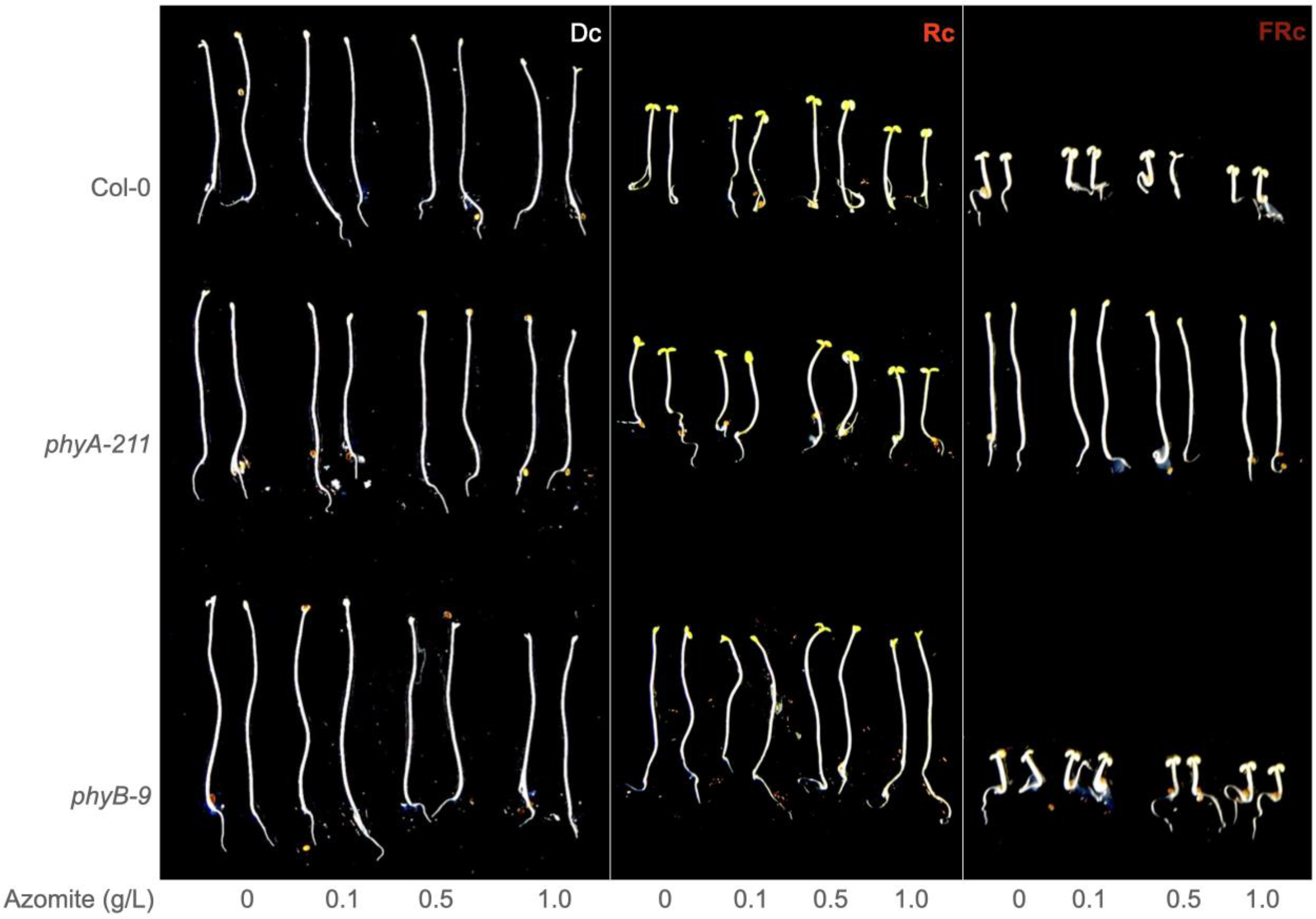
Increase in hypocotyl growth in response to Azomite is light-wavelength dependent. Representative Col-0 and phytochrome mutant (*phyA-211* and *phyB-9*) seedlings grown for 4-days in Dc, Rc, or FRc, and treated with indicated Azomite concentrations are shown.

**Fig. 5.**
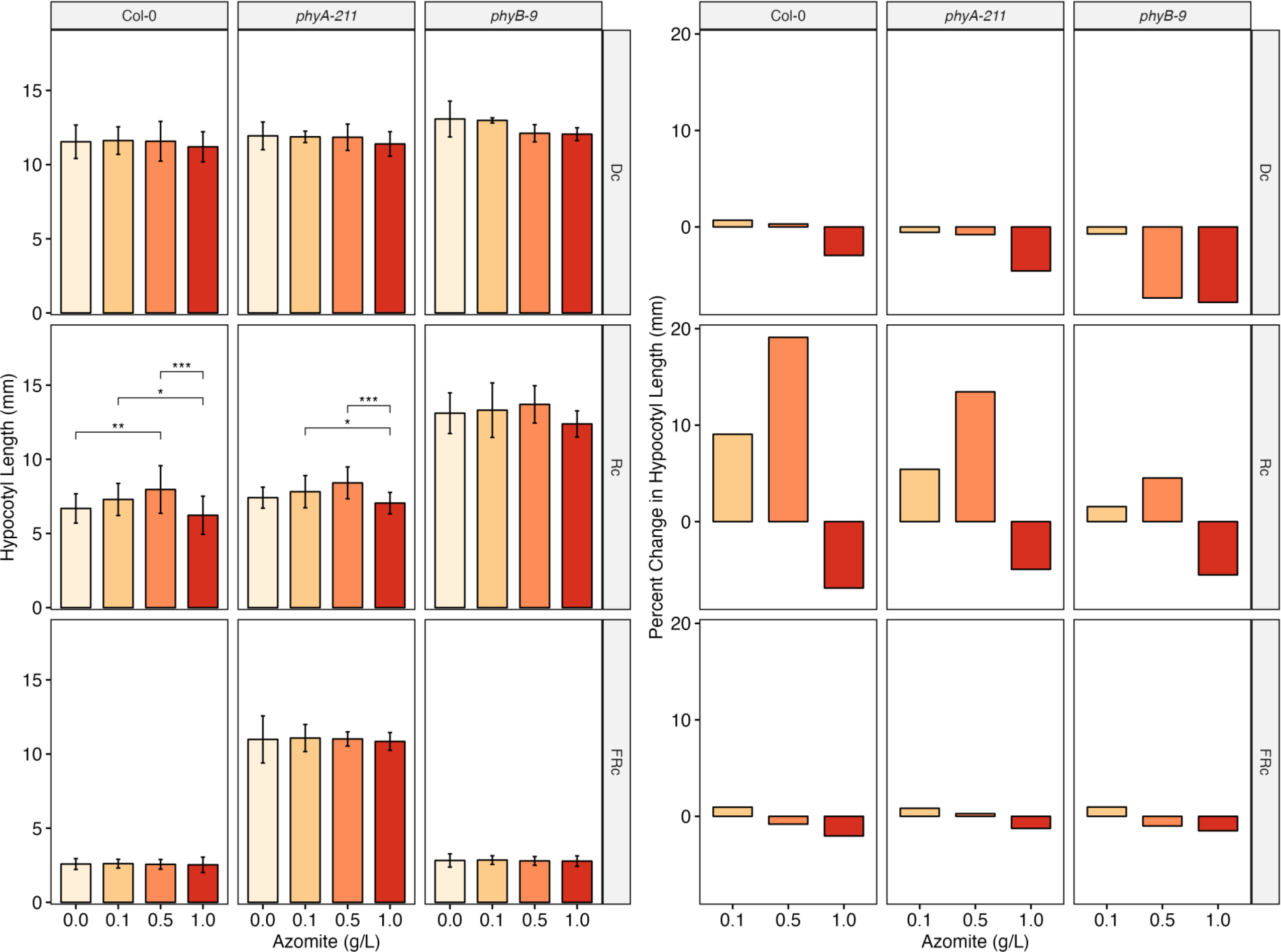
Azomite promotes hypocotyl length only under a photosynthetically active wavelength (Rc). Mean value of hypocotyl length of 4-day-old Col-0 and phytochrome mutant (*phyA-211* and *phyB-9*) seedlings grown for 4-days in Dc, Rc or FRc are plotted with corresponding Azomite treatments (left). Percent change in hypocotyl length relative to untreated (0g/L) control is shown for seedlings treated with different Azomite concentrations and light conditions (right). SE are plotted. P-values adjusted for multiple testing comparisons are represented as asterisks, with * indicating a p-value less than 0.05, ** indicating a p-value less than 0.01, *** indicating a p-value less than 0.001.

Quantitative assessment of seedling phenotypes under Rc and WLc is shown in Fig. 6. Average growth measurements of Col-0 were set to 1 for hypocotyl length and cotyledon area as previously described (Khanna et al., 2006). The plot can be divided into four quadrants representing (A) Hyposensitivity to light (top left), (B) Hypersensitivity to light (bottom right), (C) Reduced general growth (bottom left), and (D) Increased general growth (top right) (see Fig. 6). Relative mean values of hypocotyl length and cotyledon area were plotted for Col-0 grown under Rc or WLc with different amounts of Azomite as indicated, with values from 0.0 g/L Azomite treatment set to 1 (Fig. 6). This plot shows the comparative effect of lower Azomite concentrations (0.1 g/L and 0.5 g/L) on seedling growth under the two light conditions. Higher than these Azomite amounts under those conditions had a reduced growth effect (Fig. 6). Here we can draw two conclusions, one that de-etiolating seedlings exhibit dose-dependent sensitivity to volcanic ash composition, and second that Azomite activity in seedlings intersects with light-regulated development.

**Fig. 6.**
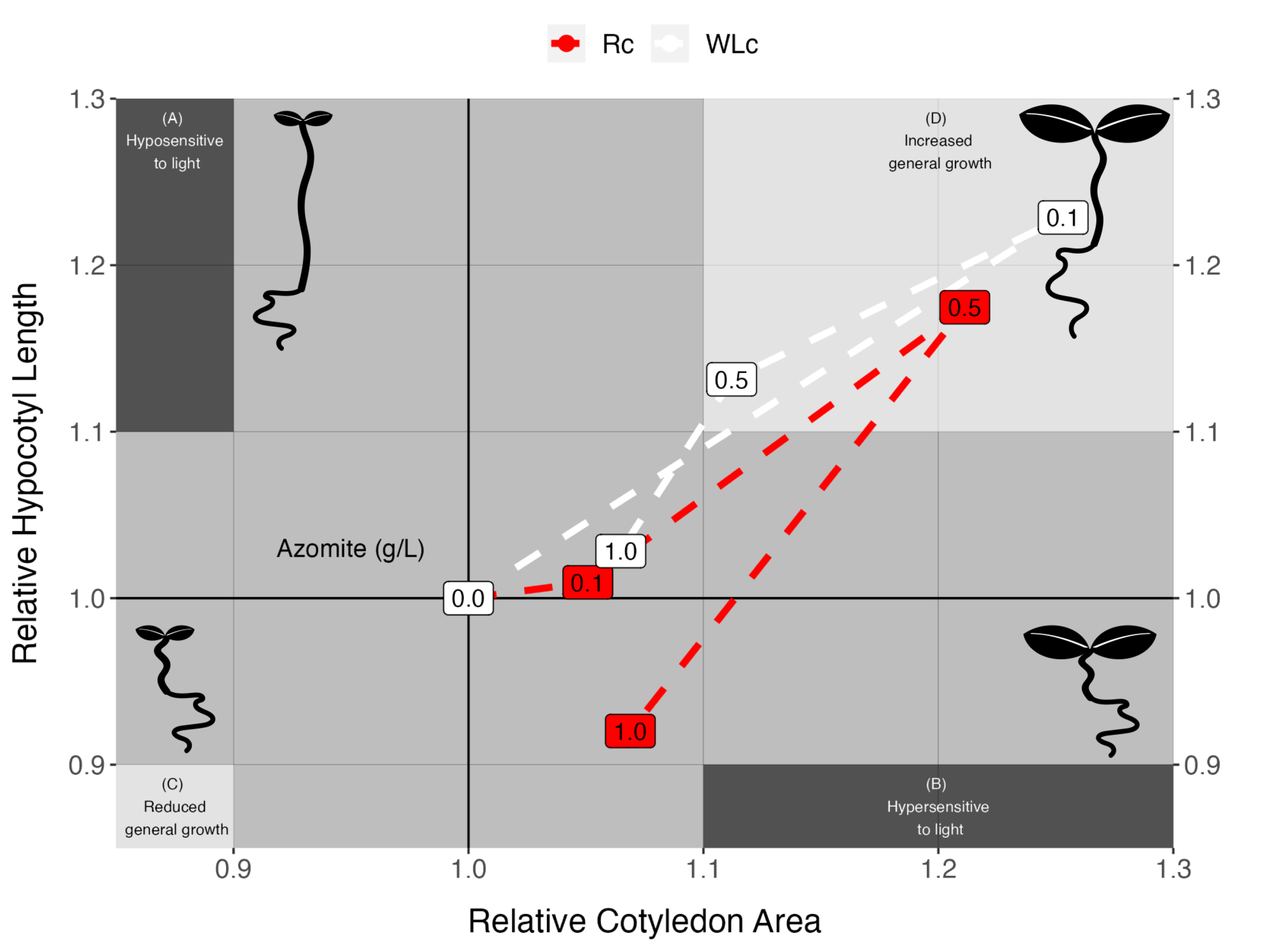
Quantitative assessment of Azomite induced seedling phenotypes. Relative mean values of hypocotyl length were plotted against relative mean values of cotyledon area, with control (0g/L) values set to unity. The quadrants represent previously described relationships of cell elongation phenotypes displayed by the two organs during deetiolation; (A) Hyposensitive, reduced light sensitivity (B) Hypersensitive, increased light sensitivity, (C) Reduced general growth, (D) Increased general growth (Khanna et al., 2006). Normal deetiolation is represented at the point of cross-section between the quadrants (0.0, control). In both Rc and WLc, increasing levels of Azomite concentration move the hypocotyl length/cotyledon area measurements closer towards increased general growth, with a concentration of 0.1 grams of Azomite maximally increasing plant growth under WLc, and 0.5 grams of Azomite maximally increasing growth under Rc. However, past these optima, general growth decreases, and in Rc at 1.0 gram of Azomite the hypocotyl length is shortened more relative to cotyledon area, moving the seedlings towards a hypersensitive response. Values plotted are from a representative experiment.

### PAR is required for Azomite induced early seedling growth

Relatively lower concentrations of Azomite increased hypocotyl and cotyledon growth under Rc and WLc. Whereas there was no significant effect of Azomite treatment on seedling growth under Dc and FRc (Figs. 2 and 5), indicating that Azomite activity required photosynthetically active radiation. These results were unexpected, and suggest that during early seedling establishment, activation of the photosynthetic circuitry precedes seedling utilization of certain micronutrients, such as those present in volcanic ash product Azomite. Data represented in Fig. 6 shows that under optimal concentrations, Azomite can promote growth processes in both organs, which is likely to involve modifications in gene expression under the conditions tested. We are in the process of testing the underlying mechanisms involved to potentially draw links between the pre-requirement of PAR-dependence for volcanic ash mediated growth stimulation during photomorphogenesis.

## Discussion

Azomite is a natural, highly mineralized complex silica ore volcanic ash deposit in Utah, of solidified volcanic igneous rock origination, comprised of a large portion of non-crystalline composite materials and characterized as a dacitic (rhyolitic) tuff breccia - not strictly a Dacite. Also described as a vitric, poorly welded tuff - meaning it is quite porous (microporous), breaks apart easily and contains in excess of 70 minerals, trace elements, and rare earth elements. Data presented here shows that the ash as a dacitic tuff breccia (DTB) used with MS salts lacking sucrose, increased growth of photomorphogenic seedlings, but not of etiolated or FRc seedlings. Early seedling growth is dependent upon environmental factors and nutrient availability. The unexpected observation that under controlled environments, Azomite effect on seedling growth required photosynthetically active radiation, suggests that one or more of Azomite components may be involved in promoting photosynthesis-related growth regulating processes during deetiolation. Normal deetiolation is characterized by concomitant inhibition of hypocotyl and stimulation of cotyledon growth (Khanna et al., 2006). In contrast, Azomite application stimulated both hypocotyl and cotyledon growth in a dose- and light-wavelength-dependent manner, and higher amounts of the volcanic ash had a growth inhibitory effect. These data provide several novel insights into the possible mechanisms underlying the beneficial effects of volcanic ash on plant growth and development.

### Volcanic ash deposits

To take advantage of the mineral nutrients present, volcanic ash-based fertilizers are applied in croplands, but dissecting the mechanisms involved in promoting plant growth is challenging due to the complex compositions and unique characteristics of each deposit. Volcanic ash soils cover nearly 124 million hectares, about 0.84% of global land surface. Soils that are formed with volcanic ash, pumice, cinders and lava develop colloidal fraction consisting of short-range-order minerals and active aluminosilicates (aluminum-humus). Weathering and mineral transformation under cool, humid climates generates andic soil properties, which is characteristic of Andisols, comprising of 60% or more of their upper layer thickness with allophane, imogolite, and ferrihydrite (Soil Survey Staff, 1999). Andisols have low bulk density, high phosphate retention, and high water-holding capacity (Shoji and Ono, 1978; Takahashi and Shoji, 2002), which leads to accumulation of organic carbon and nitrogen, making these soils one of the most productive soils in the world with long-term agricultural and environmental sustainability (Shoji and Takahashi, 2002), as evidenced by the rapid revegetation of volcanic ash deposits in areas of Mt. Usu in Japan, Mt. Pinatubo in Philippines, and Mt. St. Helens in USA.

Two patterns of revegetation have been reported; (a) establishment of seedlings on thick (1-3 m) ash deposit by wind-dispersed seeds of trees or emergence of gramineous and leguminous plants, and (b) by resprouting of buried vegetation on thin (<1 m) ash deposit (Haruki and Tsuyuzaki, 2001; Saito et al., 2002; Shoji and Takahashi, 2002). Both, Mt. Usu and Mt. Pinatubo ash deposits exhibited similar physical properties and contained abundant amounts of plant-available phosphorus (Shoji and Takahashi, 2002), but only trace amounts of organic carbon and N were present in the top layers, which were reported to increase over seven years to 0.8% organic carbon and 0.066% N with the establishment of dense gramineous communities in the vicinity of Mt. Pinatubo (Ota, 2002). Nitrogen is one of the major limiting nutrients during early revegetation of ash deposits, and symbiotic relationships of pioneer plants with diazotrophic endophytes is thought to play an important role (Shoji and Takahashi, 2002). These studies provide a glimpse into the natural progression of vegetative growth in Andisol. Nonvolcanic soils that contain weathered primary alumino-silicates may also accumulate short-range-order minerals and some of these soils qualify to be included in Andisols (Soil Survey Staff, 1999).

### Possible mechanisms of Azomite activity

At present, it is unknown how volcanic ash’s wide complex of mineral or element component/s enhance plant growth. The requirement of PAR suggests a shift in the seedling’s ability to mobilize the active ingredients. Ockenden and Lott, 1987 showed that when grown in distilled water, etiolated *Cucurbita* seedlings exported more minerals from cotyledons to the root-shoot axis than light-grown seedlings, whereas when grown under light, in soil with sufficient nutrients, the fully expanded, photosynthetic cotyledons had increased K by 5-fold and Ca by 50-fold compared to the cotyledons of dry embryos. Even in distilled water conditions, cotyledons of light grown seedlings accumulated 17% more Mg and 36% more P compared to those of dark-grown seedlings (Ockenden and Lott, 1987). Cucurbit cotyledons green early during deetiolation, and exhibited low export of Mg, which could be related to the requirement of Mg in chlorophyll synthesis (Ockenden and Lott, 1987). The DTB ash contains Mg, which could influence chlorophyll production in light grown seedlings. Studies are under way to examine this possibility. The amount of plant-available P in andisols (Truog-extractable P_2_O_5_) is similar to the amount found in fresh volcanic ash (Shoji and Takahashi, 2002), implying that majority of extractable P is plant-available. P deficiency has been shown to effect absorption of mineral nutrients and photosynthetic performance in citrus seedlings (Meng et al., 2021). A possible role of P in the Azomite ash product cannot be ruled out. During deetiolation, seedlings transition to autotrophic growth and it is possible that inclusion of Azomite minerals in the growth medium accelerated mineral mobilization under PAR.

In addition to differential nutrient remobilization between organs of etiolated and green seedlings, the mineral elements may have a regulatory function. For example, K transport into guard cells is directly related to light-regulated stomatal opening (Humble and Raschke, 1971). Both, deficiency and excess of K reduced nitrogen absorption and carbon assimilation, and inhibited growth and root development in apple dwarf rootstock seedlings, suggesting that optimal K levels are required for transport of photosynthetic products from leaves to roots and nitrate transport and assimilation (Xu et al., 2020). Import of Ca into the chloroplast is light-dependent and has been linked to regulation of photosynthesis, possibly through changes in the activities of Ca^2+^-binding proteins involved in photosynthesis electron transfer and photo-protection (Weinl et al., 2008; Hochmal et al., 2015). In maize seedlings during deetiolation, only 7% of proteins significantly changed in abundance, while there was a 26.6% change in phosphorylation status, including phosphoproteins involved in gene regulation and rate-limiting steps in light and carbon reactions of photosynthesis (Gao et al., 2020). It is conceivable that the observed PAR-dependent seedling growth promotion by Azomite was mediated through downstream regulation of photosynthesis related processes, including gene expression, post translational modification, and regulation of enzymes and transport channels. Further studies are being conducted to test these possibilities.

At the higher concentration (1 g/L) tested, Azomite reduced cotyledon expansion and hypocotyl growth in etiolated, FRc and PAR-grown seedlings (Figs. 6), implying an optimal range (>0.1 to <1.0 g/L) of beneficial Azomite activity on Arabidopsis seedlings during deetiolation. Recent greenhouse grown tomatoes have revealed similar relationships between plant growth and Azomite concentrations (Mehlferber et al., 2002a). Higher concentrations may result in an increase in growth inhibition, or it could be caused by yet unknown impact of nutrient elements like K, as described above (Xu et al., 2020).

The measured peak of Azomite’s beneficial activity shifted to lower Azomite concentrations under WLc (at 0.1 g/L), compared to Rc (0.5 g/L) (Fig. 6). This could be due to the monochromatic red (660 nm) light used compared to white light (400 – 700 nm) with a broader range of PAR. However, this observation suggests an intricate relationship between the seedling’s photosynthetic capacity and available concentration of Azomite volcanic ash in the medium. It is possible that either some growth-rate-limiting components become depleted, or relative levels of some toxic elements increase with an increase in photosynthesis processes.

In a previous study (Khanna et al., 2006) with 32 independent gene mutant lines, each corresponding to genes rapidly and robustly induced by phy-mediated light signal, and encoding putative transcription factors, signaling components, and proteins of unknown function, only one mutant line with a T-DNA insertion in a MYB family transcription factor was found to display significant parallel enhancement of both hypocotyl and cotyledon expansion, like the phenotypes presented here with optimal Azomite concentrations (Fig. 6). Subsequently, this locus was identified as *CIRCADIAN1*/*REVEILLE2* (*CIR1*/*RVE2*), belonging to a family of 11 proteins, and implicated in disrupting circadian clock-regulated gene expression (Chaudhary et al., 1999, Zhang et al., 2007). The parallel growth enhancement of both organs by Azomite is likely to be mediated through altered expression of genes in response to treatment with volcanic ash. We are investigating this possibility further. These studies will have broader implications in understanding the beneficial effects of volcanic ash and molecular processes involved in revegetation of regions impacted by volcanic deposits.

### Potential of volcanic ash fertilizers in agriculture

Application of volcanic ash to soil may create localized, transient andisol-like conditions. For example, Azomite is lightly weathered, contains short-range-order minerals with aluminosilicates, and its application in the top layer of soil may temporarily create andisol-like conditions. In previous studies with greenhouse-grown tomatoes, Azomite increased tomato production (Mehlferber et al., 2022a; 2022b), and here it was found to promote early seedling growth. These data suggest that the application of volcanic ash fertilizers, like Azomite may increase soil productivity similarly to Andislos. Higher concentrations of volcanic ash in soil can be detrimental to plant performance, and it is anticipated that the optimal concentrations will depend upon several factors, including plant size, rate of photosynthesis, other soil-borne factors, and environmental conditions. Overall, volcanic ash-based fertilizers can offer an environmentally sustainable option to promote plant growth through increased fixation of atmospheric carbon dioxide and increased C accumulation, as well as the potential to reduce or partially replace the use of chemical-based phosphorus, potassium, and other micronutrient fertilizers. Nitrogen can be supplemented through microbial activity and the inclusion of organic matter to improve soil productivity.

### Utility of photomorphogenesis bioassay to examine commercial fertilizer activity

The commercial fertilizer market was estimated to be over USD 190 billion in 2020, and it is expected to grow at a CAGR of 2.6% by 2030 (Global Market Insights, Inc., DE). There are many different fertilizers, from single components to complex additives. Common market practice is to perform field tests for product promotion with little to no insight into mechanism of activity or the impact on the soil and the environment. The photomorphogenesis bioassay presented here offers an easy entry path to determining the possible activities, particularly for commercial fertilizers that can be applied to deetiolating seedlings without impacting the bioassay performance. Deetiolation in Arabidopsis seedlings is highly sensitive to alterations in growth conditions, and it is one of the most studied phenomena at genetic, molecular and biochemical levels, providing access to established knowledge for data interpretation and tools, such as mutant lines for further investigation.

## Materials and Methods

### Plant materials and growth conditions

*Arabidopsis thaliana* Columbia ecotype (Col-0), *phyA-211*, or *phyB-9* seeds were sterilized (Khanna et al., 2006) and plated on half-strength MS (Murashige & Skoog Basal Salt Mixture, PhytoTechnology Laboratories) salts, buffered with 1 g/L MES (2-(N-Morpholino) Ethane Sulfonic Acid, Research Organics, Inc.) at pH 5.8 with KOH, plus 0.7% Agar Noble (Difco laboratories). The media did not contain any added sugars. Azomite Ultrafine (Azomite Mineral Products, Inc.) was added at various concentrations; 0 mg/mL, 0.1mg/mL, 0.5 mg/mL, and 1.0 mg/mL prior to autoclaving, and Azomite was maintained as a uniform suspension in the media by shaking while pouring the plates. The experiments were repeated 3 to 5 times consistently by stratifying the plates for 4 days at 4°C; germination was induced by 3 hours of white light treatment in a growth chamber at 21°C, followed by continuous placement in the growth chamber in darkness for 21 hours. The plates were then transferred to respective light treatments in various growth chambers for 4-days for seedling growth either in Dc, WLc (5 μmol.m^−2^.sec^−1^), Rc (660 nm, 7 μmol.m^−2^.sec^−1^), FRc (740 nm, 2 μmol.m^−2^.sec^−1^), as indicated. Fluence rates were monitored using a spectroradiometer (model L1–1800; LICOR).

### Measurements and statistical analyses

Hypocotyl length and cotyledon area measurements were performed 96-hours after germination using a digital camera (Canon EOS DSLR) and Image J software (National Institutes of Health) and analyzed using Excel (Microsoft). Mean values were calculated to determine relative differences between untreated and treated seedlings. For all statistical tests each group was analyzed separately (split either by light, or by light and mutant line) with an ANOVA (using anova test in r) estimating the impact of Azomite concentration under those conditions on Hypocotyl length or Cotyledon area. Significant effects were further probed by performing a Tukey posthoc test (using tukey hsd in r) where p values were adjusted for multiple testing comparisons. Data presented are representative of minimum of three replicates, each with 10 to 35 seedlings.

### Azomite Volcanic ash composition

Azomite (A to Z Of Minerals Including Trace Elements, AZOMITE^®^ Mineral Products, Inc.) is surface mined from naturally deposited volcanic ash (Nephi, UT). In composition, Azomite is a lightly weathered DTB, it is classified as a hydrated calcium sodium aluminosilicate and listed as an anticaking agent (United States Code of Federal Regulations, 21 CFR 582.2729). It is a natural, highly mineralized complex silica ore, with on average over 70 minerals, trace elements, 529.7 ppm of total Rare Earth Elements, including lanthanides (Chemical analysis, AZOMITE^®^ Mineral Products, Inc.). Azomite is a mixture of different substances present in pulverized bits of rock, crystalized minerals, and volcanic glass (Dr. Barry Bickmore, Brigham Young University, *personal communication*); it is produced as a product in different grades from Granulated to Micronized (AZOMITE^®^ Mineral Products, Inc.). For the work presented here, some of the known agriculturally important elements found in Azomite formulations are potassium, phosphorus, calcium, sodium, iron, magnesium, and manganese, along with trace amounts of zinc, copper, molybdenum, selenium. Based upon multiple sample analysis, the manufacturer provides a guaranteed analysis of (0-0-0.2 of N-P-K), along with 1.8% Ca, 0.5% Mg, and 0.1% of Cl and Na. Potassium is present as soluble potash (K_2_O).

## Abbreviations

PAR: Photosynthetically Active Radiation
Dc: continuous Darkness
WLc: continuous White light
Rc: continuous Red light
FRc: continuous Far-Red light
DTB: dacitic (rhyolitic) tuff breccia

## Acknowledgements and Funding

The authors declare that this work was funded and performed through i-Cultiver, Inc., which independently provides consultation services to soil amendment manufacturers; this includes AZOMITE^®^ Mineral Products, Inc., which manufactures the Azomite product. This funding provided for research costs associated with this study. It did not influence the factual reporting of findings herein. Robert and Alexis were supported by i-Cultiver’s Biotechnology Education and Specialized Training (B.E.S.T.) internship program in collaboration with Dr. Katie Krolikowski, (Contra Costa Community College, Richmond, CA) and Dr. Zhiyong Wang (Carnegie Institution for Science at Stanford University, CA). The authors thank Pratigiya Khatiwada, another B.E.S.T. program intern who joined towards the completion of the current study and assisted with preparing media plates and plating seeds.

## Author Contributions

R.K. conceived and designed the study; K.M. and R.K. conducted the experiments, obtained seedling images and data with support in plating and measurements from R.R. and A.O.; E.M. performed statistical analysis and produced graphical figures; J.F. provided product-specific details in the study design. K.M. and R.K. drafted the manuscript supplemented with technical and editorial comments from all authors.

## Disclaimer

Mention of trade names or commercial products in this publication is solely for the purpose of providing specific information and does not imply recommendation or endorsement by the U.S. Department of Agriculture. USDA is an equal opportunity provider and employer.

